# The MarR Family Regulator BmrR is involved in Bile Tolerance of *Bifidobacterium longum* BBMN68 via Controlling the Expression of an ABC-Transporter

**DOI:** 10.1101/310508

**Authors:** Qi Xu, Zhengyuan Zhai, Haoran An, Yang Yang, Jia Yin, Guohong Wang, Fazheng Ren, Yanling Hao

**Affiliations:** The Innovation Centre of Food Nutrition and Human Health (Beijing), College of Food Science and Nutritional Engineering, China Agricultural University, Beijing, China; Key Laboratory of Functional Dairy, Co-constructed by Ministry of Education and Beijing Municipality, Beijing, China; Department of Food Science & Technology, University of California, Davis, California, USA; Center for Infectious Disease Research, Tsinghua-Peking Joint Center for Life Science, School of Medicine, Tsinghua University, Beijing, China

**Keywords:** *B. longum* BBMN68, bile stress, MarR-type regulator, BmrR, ABC-transporter

## Abstract

In order to colonize the human gastrointestinal tract and exert their beneficial effects, bifidobacteria must effectively cope with the toxic bile salts in the intestine, but the molecular mechanism underlying bile tolerance is poorly understood. In this study, heterologous expression of a MarR family transcriptional regulator BmrR significantly reduced ox-bile resistance of *Lactococcus lactis* NZ9000, suggesting that it might play a role in bile stress response. *In silico* analysis combined with RT-PCR assay demonstrated that *bmrR* was co-transcribed with *bmrA* and *bmrB,* which encoded multidrug resistance (MDR) ABC transporters. Promoter prediction and EMSA assay revealed that BmrR could autoregulate the *bmrRAB* operon by binding to *bmr* box (ATTGTTG-6nt-CAACAAT) in the promoter region. Moreover, heterologous expression of *bmrA* and *bmrB* in *L. lactis* showed 20.77-fold higher tolerance to 0.10% ox-bile compared to wild type strain. In addition, ox-bile could disrupt the DNA binding activity of BmrR as a ligand. Taken together, our findings indicate that *bmrRAB* operon is autoregulated by transcriptional regulator BmrR and ox-bile serves as an inducer to activate the bile efflux transporter BmrAB in response to bile stress in *B. longum* BBMN68.

**Importance:** Bifidobacteria are natural inhabitants of the human intestinal tract. Some bifidobacterial strains are used as probiotics in fermented dairy production because of their health-promoting effects. Following consumption, bifidobacteria finally colonize the lower intestinal tract where the concentration of bile salts remains nearly 0.05% to 2.0%. Bile salts as detergent-like antimicrobial compounds can cause disruption of the cellular membrane, protein misfolding and DNA damage. Therefore, tolerance to physiological bile stress is indeed essential for bifidobacteria to survive and exert the probiotic effects in gastrointestinal tract. In *B. longum* BBMN68, the MarR-type regulator BmrR was involved in bile stress response by auto-regulating *bmrRAB* operon and ox-bile as an inducer could increase the expression of BmrAB transporter to enhance the bile tolerance of BBMN68.This is the first report about functional analysis of *bmrRAB* operon in bile stress response, which will provide new insight into bile tolerance mechanisms in *Bifidobacterium* and other bacteria.

## Introduction

Bifidobacteria are natural inhabitants of the human gastrointestinal tract (GIT), constituting up to approximately 91% of the total gut microbiome during early stages of life (1). Some bifidobacteria are considered as probiotics, and used as active ingredients in functional dairy-based products (2). The health benefits are exerted mainly through inhibiting pathogens, preventing diarrhoea, stimulating the immune response and reducing serum cholesterol levels (3). Upon ingestion, bifidobacteria inevitably have to cope with several stress conditions, such as the low pH in the stomach and bile salts in the intestine (4, 5). As detergent-like biological substances with strong antimicrobial activities, bile salts can disrupt the lipid bilayer structure of cellular membranes, induce protein misfolding and cause DNA damage (6). Bifidobacteria have been reported to develop tolerance response to bile stress, but the comprehensive mechanism of bile resistance remains elusive.

Among the bile resistance mechanisms employed by bifidobacteria, bile salt hydrolysis (BSH) and bile efflux transporter are well documented. Bile salt hydrolases (BSHs) are responsible for deconjugation of glycine- or taurine-conjugated bile salts, therefore decreasing the toxicity of conjugated bile salts (7). The bile efflux system is mediated by a multidrug resistance (MDR) transporter located on the cell membrane, such as Ctr of *B. longum* NCIMB 702259^T^ (8), BetA in *B. longum* NCC2705 (9) and BbmAB in *B. breve* UCC2003 (10). Several studies have shown that bifidobacteria modulated the cell envelope including fatty acid composition and membrane proteins to decrease membrane permeability in response to bile salts (11, 12). In addition, a hemolysin-like protein TlyC1 functions as a barrier to protect the strain from bile toxicity and provides resistance to sodium taurocholate and sodium taurodeoxycholate in *B. longum* BBMN68 (13). Two-component system *senX3-regX3* was reported to promote the expression of the *pstS* gene to maintain a high-level of Pi uptake and produce more ATP to resist bile stress in *B. longum* BBMN68 (14).

*B. longum* BBMN68 was isolated from healthy centenarians in Bama longevity villages of Guangxi province in China, which may enhance innate and adaptive immunity, alleviate allergic response and improve intestinal function in mice (15, 16). In our study, RNA-Seq transcriptomic analysis showed that the BBMN68_1796 gene encoding MarR-type transcriptional regulator was 1.85-fold up-regulated under bile stress in BBMN68 (unpublished). It has been reported that the MarR family transcriptional regulators are involved in the regulation in response to diverse environmental signals, such as synthesis of virulence factors and antibiotic stress (17, 18). Martin *et al.* found that transcription of multiple antibiotic resistance (marORAB) operons was repressed by the MarR protein (19). Furthermore, another MarR type repressor EmrR has been reported to control the EmrAB efflux transporter in *E. coli* (20). In the present work, we investigated the regulatory mechanism of protein BBMN68_1796 designated as BmrR *(Bifidobacterium* multidrug resistance regulator) in ox-bile stress response in *B. longum* BBMN68. The data suggests that BmrR autorepresses the transcription of *bmrR* operon and ox-bile serves as ligand of BmrR to attenuate this binding to enhance the expression of efflux transporter genes to export ox-bile in *B. longum* BBMN68.

## Results

### Heterologous expression of *bmrR* in *L. lactis* NZ9000 increases its sensitivity to bile stress

DNA sequencing showed that the length of the amplified gene *bmrR* was 534 bp, which was 100% homologous to the *bmrR* gene from *B. longum* BBMN68 (BBMN68_1796; GenBank Accession No. NC_014656.1). SDS-PAGE analysis revealed that the production of an expected 20 kDa protein in *L. lactis* BmrR after nisin induction (Figure 1A, lane 4), indicating the successful expression of *bmrR* in *L. lactis* NZ9000. Then, the recombinant strains were incubated with 0.10% wt/vol ox-bile for 1 h, and the survival of *L. lactis* BmrR was 57-fold lower than that of *L. lactis* NZCK (P<0.0001, Figure 1B). These results showed that the heterologous expression of *bmrR* in *L. lactis* NZ9000 significantly reduced its resistance to ox-bile, indicating that BmrR played a critical role in bile stress response.

**Figure 1.**
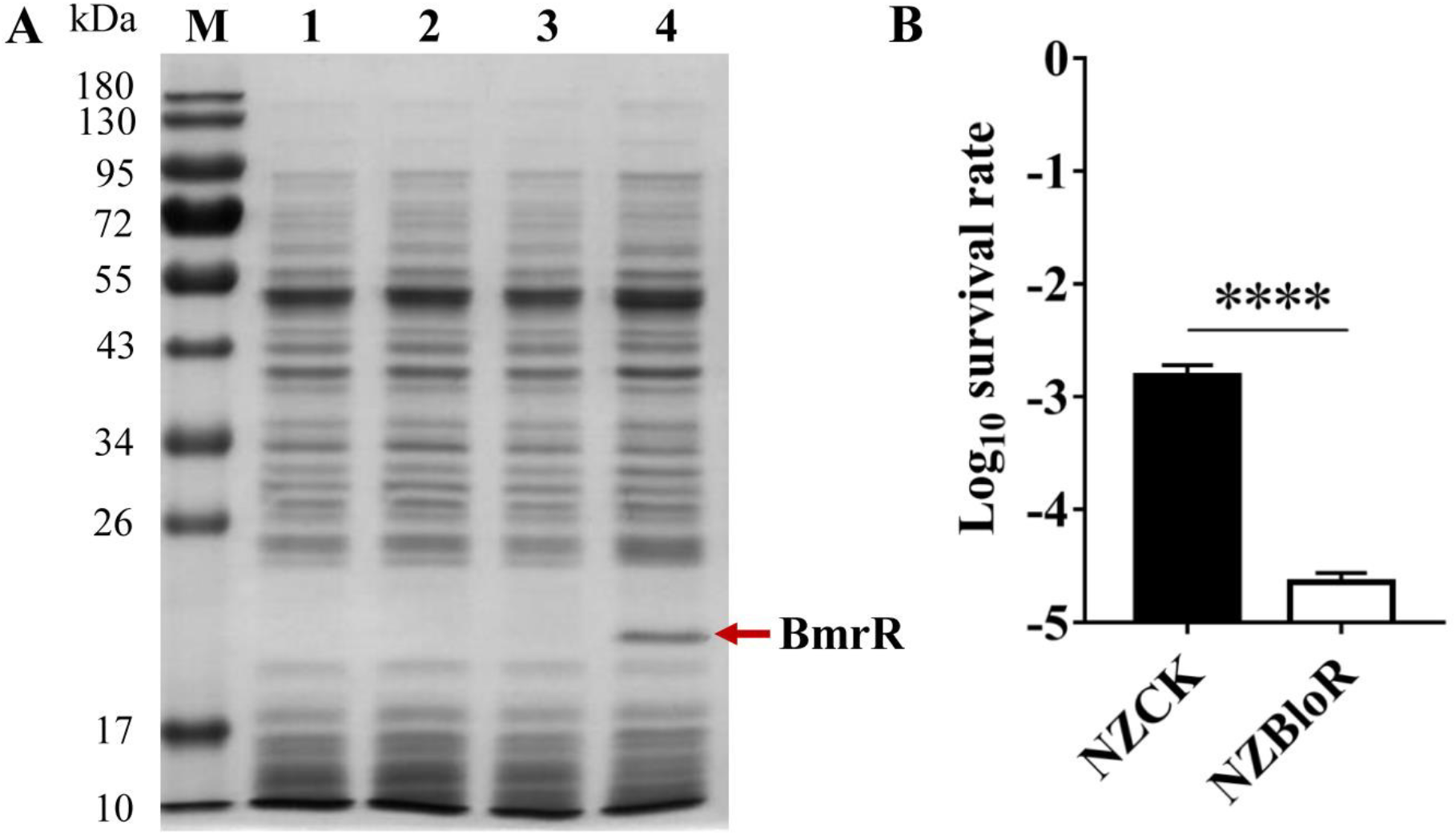
Heterologous expression of BmrR with nisin induction detected by SDS-PAGE and the survival of *L. lactis* BmrR and *L. lactis* NZCK after ox-bile challenge. (A) Soluble extracts were analyzed on 12% denaturing SDS-PAGE. Lane M, Dual color pre-stained broad molecular weight protein marker (10-180 kDa). Lane 1, *L. lactis* NZCK without nisin induction; lane 2, *L. lactis* BmrR without nisin induction; lane3, *L. lactis* NZCK with nisin (10 ng · ml^−1^) induction; lane 4, *L. lactis* BmrR with nisin (10 ng · ml^−1^) induction. Red arrow indicates the overexpressed BmrR. (B) Survival rate is calculated as the ratio of the number of colonies obtained on GM17 plates after and before ox-bile treatment. Data are reported as mean±SD from at least three independent experiments and analyzed by an unpaired, two-tailed Student t-test. ****, *P*<0.0001.

### Bioinformatics analysis revealed that BmrR was a MarR family regulator

In *B. longum* BBMN68, BmrR is annotated as a putative MarR family regulator. Although the amino acid sequence similarity between MarR family members was usually less than 25% as described previously (21), secondary structure prediction of BmrR revealed a high structural homology with the MarR family members used in the alignment (Figure 2). The core of the domain consists of three α-helices (α1, α2 and α3) and two antiparallel beta sheets (β2 and β3), which is stabilized in part by a short beta strand (β1), creating a three-stranded beta sheet (17, 22). These MarR proteins have been described as winged helix proteins that bind directly to the DNA to control a wide range of biological processes in both bacteria and archaea (23, 24). The region spanning amino acids 61-121 in MarR are required for its DNA binding activity (25). The DNA binding motif is composed of β1–α3–α4–β2–β3, which adopts the winged-helix fold. Helices α1, α5 and α6 are involved in dimerization (23, 26), which indicates that they could bind to the promoter region of their target genes as dimers leading to either transcriptional repression and/or activation (17).

**Figure 2.**
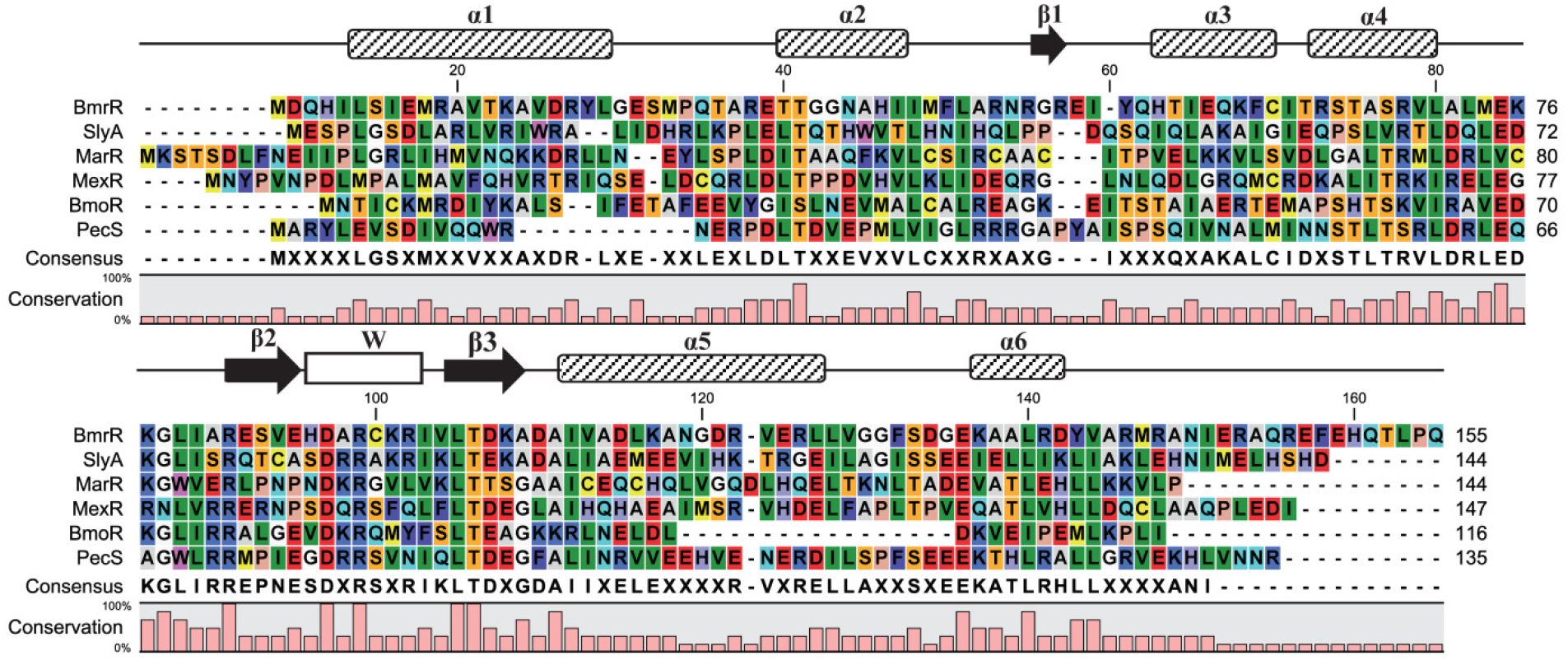
Multiple sequence alignment of BmrR with other MarR family regulators was generated with ClustalX and visualized in CLC Sequence Viewer 7.8.1. Numbering is according to the entire alignment. The proteins used for the alignment were from the following organisms: SlyA, *Salmonella enterica* sv. Typhimurium; MarR, *E. coli*; MexR, *Pseudomonas aeruginosa*; BmoR, *Bacteroides fragilis*; PecS, *Dickeya dadantii.* The prediction of BmrR secondary structure was based on Psipred and NetSurfP results and structures are illustrated with boxes (α-helices), arrows (β-sheets) and lines (coils). The wing region (W) is indicated by white box.

### *In silico* analysis of the *bmrR* gene and determination of *bmrRAB* operon

We noticed that the start codon of *bmrR, bmrA* and *bmrB* genes were overlapped with the stop codon of the proceeding gene. A putative promoter sequence was found 64 bp upstream of the potential *bmrR* start codon by the online promoter prediction tools NNPP and BPROM (27, 28), but no other promoter was predicted upstream the *bmrA* or *bmrB* gene. The first gene *bmrR* possessed a putative ribosome-binding site 8 bp (TGGTAC) upstream of its start codon, while the third gene *bmrB* was followed by a transcription terminator-like sequence (Figure 2A). Based on these observations, we hypothesized that these three genes were co-transcribed in the same cluster. RT-PCR assay with cDNA as template further confirmed that genes from *bmrR* to *bmrB* formed a polycistronic operon, designated as *bmrRAB* (Figure 3B). MarR family regulators were reported to bind recognizable palindromic sequences within the promoter region upstream the target genes (29). Bioinformatics analysis revealed that an IR sequence (ATTGTTG-6nt-CAACAAT) was also found within the *bmrRAB* promoter in BBMN68.

**Figure 3.**
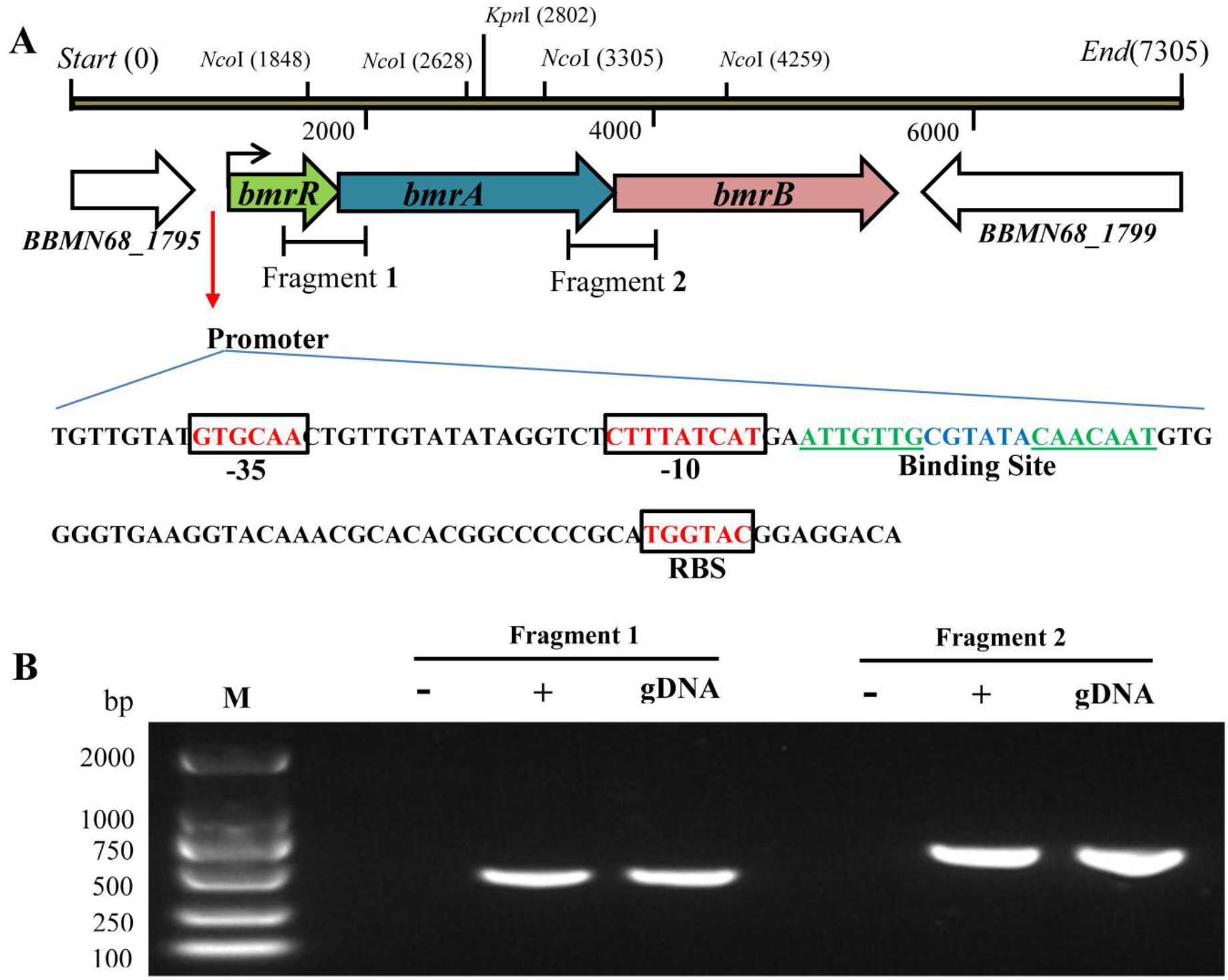
*In silico* analysis and RT-PCR assays to verify the co-transcription of *bmrR* to *bmrB.* (A) Linear map of *bmrR, bmrA* and *bmrB* with the genomic DNA flanking these genes in BBMN68. (B) Sequence analysis of the promoter region upstream *bmrR* gene. Putative -35, -10 sequence and ribosome-binding site (RBS) are enclosed in the box. The putative binding site is shown in italics. (C) “gDNA” means the genomic DNA of wide-type BBMN68; “+” and “-” indicated the cDNA and RNA used as the template for PCR amplification. “M” means the DNA marker.

### Identification of the DNA-binding specificity of BmrR by EMSA

In order to further confirm DNA-binding specificity of BmrR with its promoter, a 69 bp DNA probe was synthesized and labeled by biotin at 3’end for EMSA. The BmrR with a C-terminal His tag was expressed in *L. lactis* NZ9000 and purified by affinity chromatography. SDS-PAGE revealed a single protein band with a molecular mass of approximate 20 kDa, indicating that the recombinant BmrRHis was successfully expressed and purified for subsequent EMSAs (Figure 4A, lane 4). The EMSA results indicated that the purified BmrRHis bounded to biotin-labeled *bmrR* probe and retarded its mobility (Figure 4C, lane 2). Moreover, the quantity of DNA-protein binding bands was enhanced with an increasing concentration of BmrR (Figure 4C, lane 2 to lane 4). BmrRHis could not bound to either mutated probe up^−^ or probe down^−^ (Figure 4D), indicating the palindromic sequence (ATTGTTG-6nt-CAACAAT), designated as *bmr* box in *bmrRAB* promoter region was essential for BmrR binding. These findings indicated that BmrR could specifically bind to *bmr* box upstream the *bmrRAB* operon.

**Figure 4.**
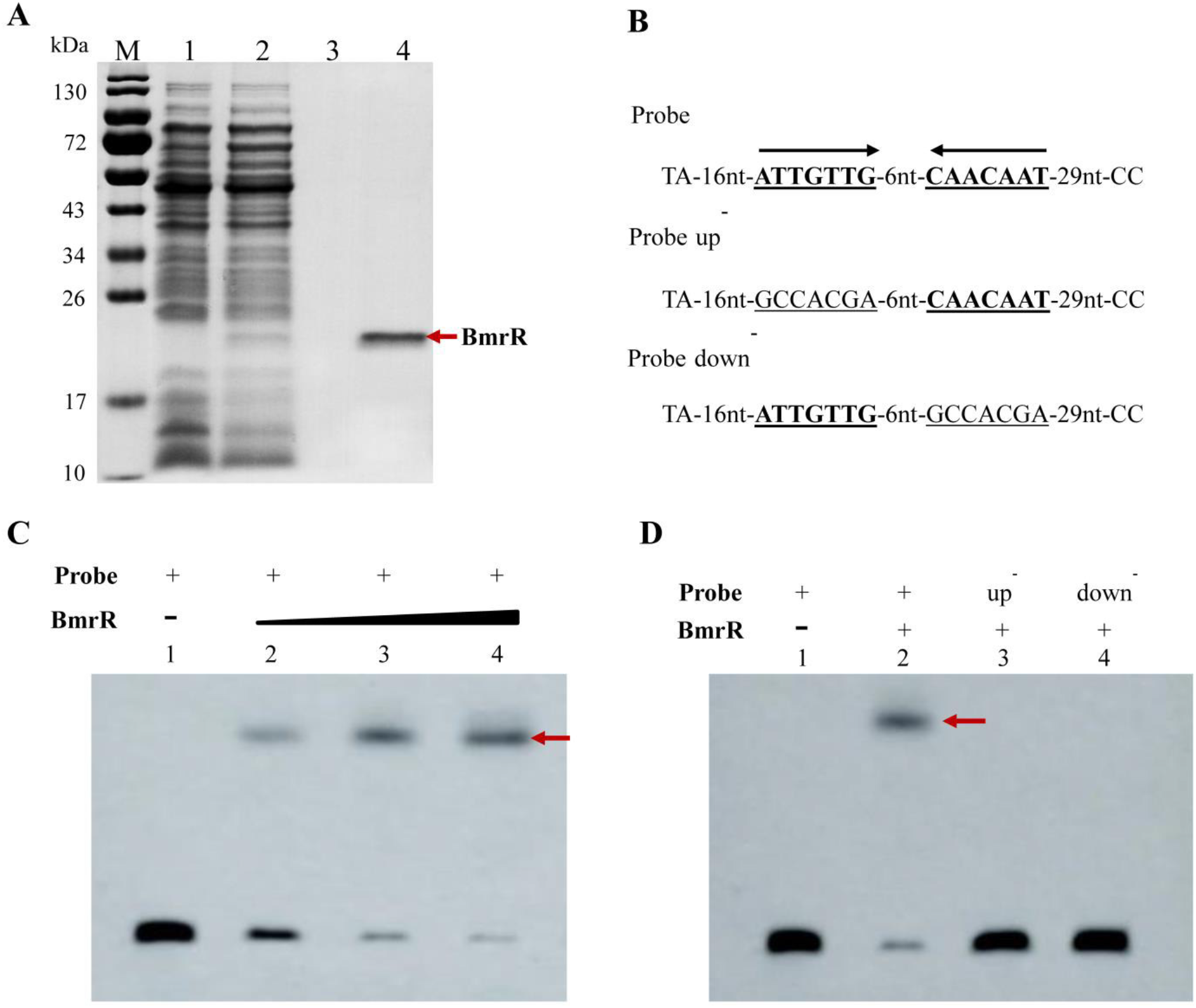
SDS-PAGE analysis of the purified BmrRHis and specific binding of BmrRHis to its own promoter. (A) Lane 1 and 2, *L. lactis* NZCK and *L. lactis* BmrRHis with 10 ng · ml^−1^ nisin induction; Lane 3, protein sample from NZCK after purification; Lane 4, purified recombinant BmrRHis from *L. lactis* BmrRHis. (B) DNA probes containing an intact palindromic sequence in the BmrR binding site or mutated sequence. (C) Lane 1, 20 fmol labelled probes alone. Lane 2 to lane 4, 20 fmol probes and 10, 20 and 30 ng · μl^−1^ BmrRHis, respectively. (D) Lane 1, 20 fmol labelled probes alone. Lane 2 to lane 4, 10 ng · μl^−1^ BmrRHis with 20 fmol probes, 20 fmol probe up^−^ and probe down^−^, respectively.

Heterodimer ABC-transporter BmrAB was involved in ox-bile tolerance In order to determine whether the ABC-transporter BmrAB was involved in the ox-bile resistance, *bmrA, bmrB* and *bmrAB* were amplified and cloned into pNZ8147 vector. The recombinant plasmids were verified by DNA sequencing and then transformed into heterologous host *L. lactis* NZ9000, resulting in *L. lactis* BmrA, *L. lactis* BmrB and *L. lactis* BmrAB, respectively. Survival assay showed that there was no significant difference between the recombinant strain *L. lactis* BmrA and control strain *L. lactis* NZCK under bile stress (P>0.05, Figure 5), but the survival rate of *L. lactis* BmrB was 16-fold lower than that of *L. lactis* NZCK in presence of 0.10% wt/vol ox-bile. It is noteworthy that the survival rate of BmrA and BmrB co-expressed strain *L. lactis* BmrAB was significantly increased, which was 20.77-fold higher than that of the control in GM17 supplemented with 0.10% wt/vol ox-bile (P<0.05, Figure 5). These results indicated that BmrA and BmrB together can enhance the bile resistance of host strain probably by forming a heterodimer ABC-transporter to pump out the intracellular bile.

**Figure 5.**
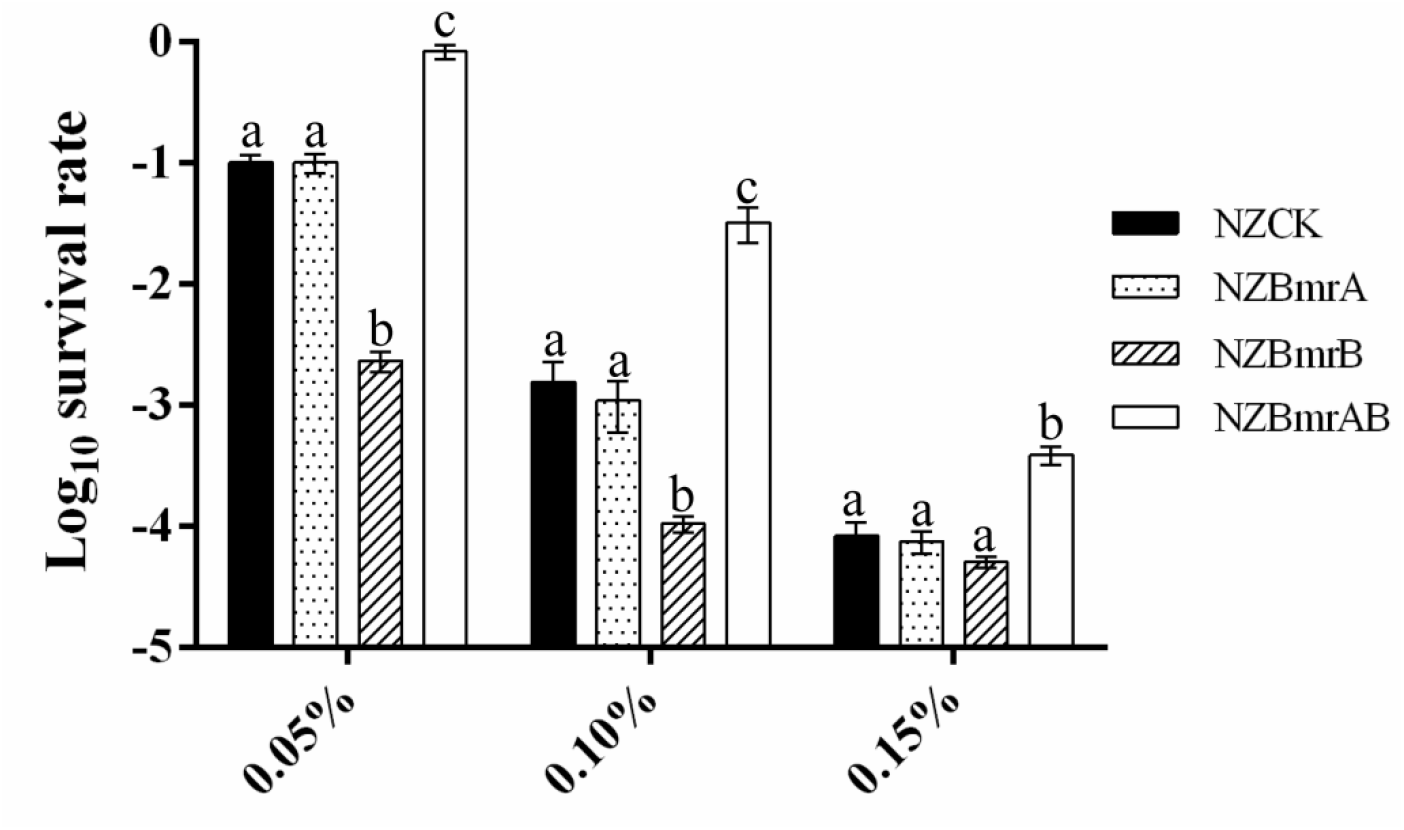
The heterologous expression of gene *bmrA, bmrB* and *bmrAB* in *L. lactis* affects the survival of host strain after bile stress. Survival rate is calculated as the ratio of the number of colonies obtained on GM17 plates after and before ox-bile treatment. Data are reported as mean±SD from at least three independent experiments and analyzed by an unpaired, two-tailed Student t-test. Bars with different letters are statistically significant *(P* < 0.05).

### BmrR dissociates from DNA in the presence of ox-bile

The DNA binding activity of some transcriptional regulators from MarR family was reported to be affected by specific ligands, which dissociate the regulator from DNA with a consequent modulation of gene expression (30). In our study, the mRNA level of *bmrR* was upregulated 1.85-fold under ox-bile stress in BBMN68 (unpublished). Therefore, we hypothesized that ox-bile might be a ligand of BmrR and affect the interaction between BmrR and its binding site. To verify this hypothesis, different concentrations of ox-bile were applied to EMSA reactions. The addition of 0.15% wt/vol ox-bile led to complete dissociation of BmrR from its target DNA (Figure 6, Lane 5). These results indicated that ox-bile was an effector for BmrR and could disrupt the DNA binding activity of BmrR in BBMN68.

**Figure 6.**
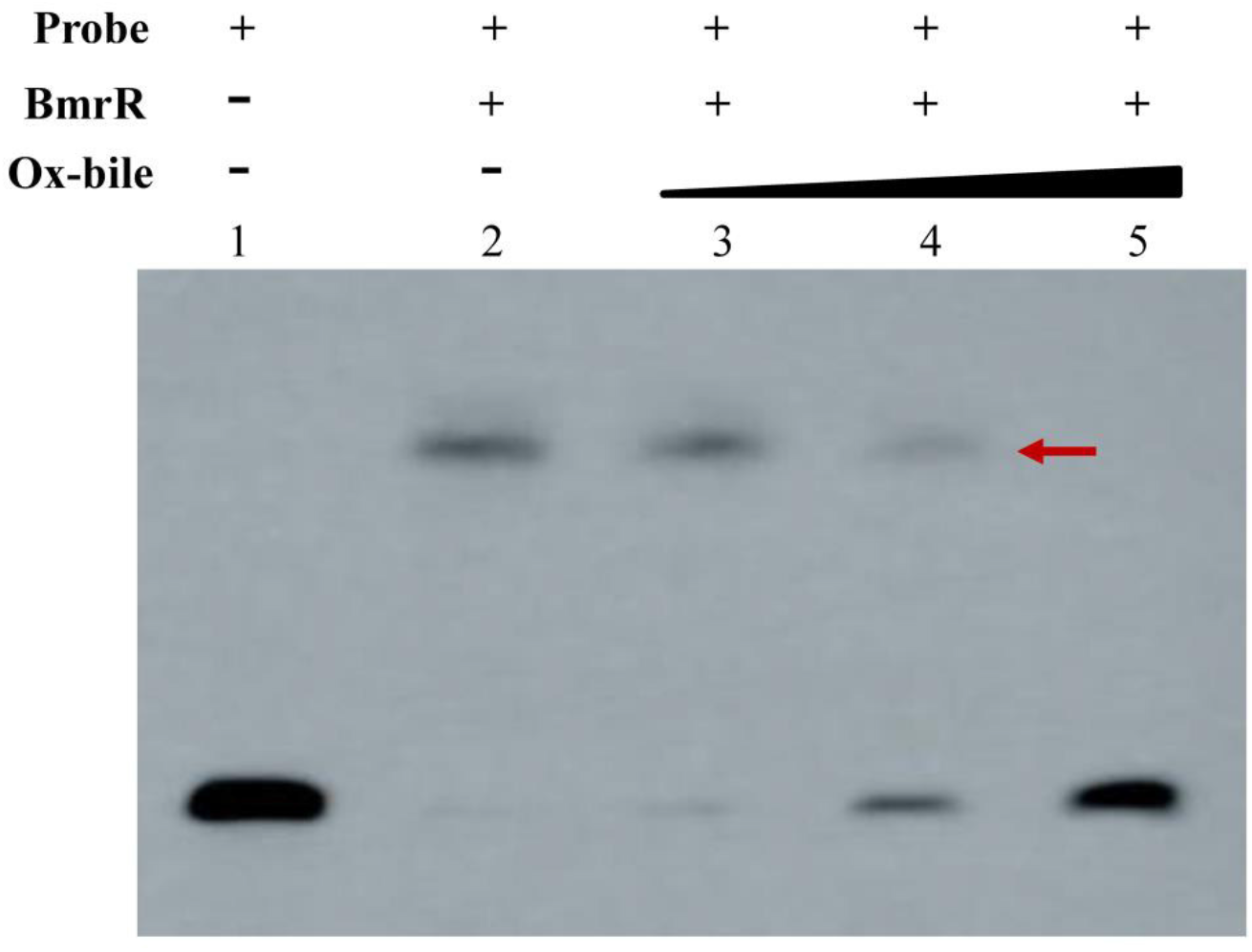
Effect of ox-bile on the DNA binding activity of BmrR. EMSA was performed with 20 fmol probe and 10 ng · μl^−1^ purified BmrRHis in the presence of 0.05%, 0.10% and 0.15% wt/vol ox-bile, respectively.

## Discussion

In bacteria, multiple antibiotic resistance regulator (MarR) family proteins constitute a diverse group of transcriptional regulators that modulate the expression of genes encoding proteins involved in metabolic pathways, stress responses, virulence and degradation or efflux of harmful chemicals (22). Some MarR family transcription factors involved in bile stress response have been identified in multiple bacterial species, including *Salmonella Typhimurium*, *L. lactis* and *Enterococcus faecalis* (31–33). In *Salmonella Typhimurium, marRAB* was activated in the presence of bile and the *marRAB* mutant showed more sensitive to bile stress than the wild-type strain (31). SlyA is a MarR family transcriptional regulator identified in *Enterococcus faecalis* and the growth of *slyA* mutant strain was significantly impaired in the presence of bile salts (33). In this study, heterologous expression of *bmrR* gene in *L. lactis* NZ9000 decreased the bile tolerance of host strain, suggesting that it might play a role in bile stress response. The *bmrR* gene was co-transcribed with *bmrA* and *bmrB*, which encoded the multidrug resistance (MDR) ABC transporters. This ABC transporter BmrAB was observed to enhance the bile tolerance of host strain, when *bmrAB* gene was expressed in *L. lactis* NZ9000. Therefore, we supposed that the *bmrRAB* operon in bifidobacteria played a critical role in enhancing the resistance to bile stress.

The *bmrA* and *bmrB* gene were predicted to encode 652 and 671 amino acid protein as putative ABC transporters by a database enquiry (BLASTP). The hydropathic profile analysis demonstrated that both proteins possessed a transmembrane domain with six putative helices, followed by cytoplasmically localized NBD domain with a putative ATP-binding domain and the ABC signature sequence (34, 35). The ATP-hydrolyzing domains are characterized by two short sequence motifs in their primary structure (‘Walker’ site A and ‘Walker’ site B) that constitute a nucleotide binding fold (Figure S1). These analyses suggested that BmrA and BmrB might also serve as ABC half-transporter. The ABC transporter utilized the free energy of ATP hydrolysis to drive substrate transport across lipid bilayer. It has been proved that several prokaryotic ABC transporters act as dimers. Homodimeric ABC transporters have been experimentally identified, such as LmrA in *L. lactis* (36) and MsbA in *E. coli* (37). Meanwhile, heterodimeric ABC transporters are also found in some species, such as LmrCD in *L. lactis* (38) and BbmAB in *B. breve* UCC2003 (10). BbmA and BbmB were further reported to be induced by 3.21±1.3 fold and 5.00±0.9 fold in the presence of bile salts, respectively (9). In agreement with these MDR transporters, BmrA and BmrB were found to be 2.33 fold and 2.09 fold up-regulated by ox-bile in *B. longum* BBMN68 (unpublished). In this study, the co-expression of BmrA and BmrB can enhance the bile resistance of host strain, suggesting that BmrA and BmrB formed a heterodimer ABC-transporter to pump out the intracellular bile in *B. longum* BBMN68.

In the present study, we observed that the MarR family regulator BmrR could interact with the promoter region of *bmrRAB* operon to regulate the transcription of these three genes. In addition, the 3D structure of BmrR was generated using SWISS MODEL server (http://www.expasy.ch/swissmod/SWISS-MODEL.html), indicating BmrR is able to form a homodimer like other MarR proteins (Figure S2). Generally, MarR family regulators were reported to bind recognizable palindromic sequences within the promoter region, resulting in attenuation of gene expression by sterically hindering the binding of RNA polymerase to the promoter. In addition, the MarR family transcription factors can also respond to a variety of effector molecules (17, 22). When the ligand binds to MarR family transcription factor, DNA binding is attenuated, resulting in de-repression of transcription (17). In this study, we observed that the formation of BmrR-DNA complex was impaired in the presence of ox-bile (Figure 6). Based on these results, we proposed a bile sensing and adaptive regulation model of *bmrRAB* operon in *B. longum* (Figure 7). Under the normal growth condition, BmrR binds to *bmr* box in the *bmrRAB* promoter region and prevents transcription of *bmrRAB* operon. When ox-bile enters the cell, BmrR interacts with ox-bile and then causes significant conformational change in the DNA-binding domains, resulting in release of the BmrR repressor from *bmrRAB* promoter. This modification will lead to the transcription of BmrAB ABC-transporters. Newly synthesized BmrAB will be embedded in the membrane and mediate the efflux of the ox-bile from the cell. To our knowledge, this is the first report about functional analysis of *bmrRAB* operon in bile stress response, which is of great importance for exploring novel bile tolerance mechanisms in *Bifidobacterium* and other bacteria.

**Figure 7.**
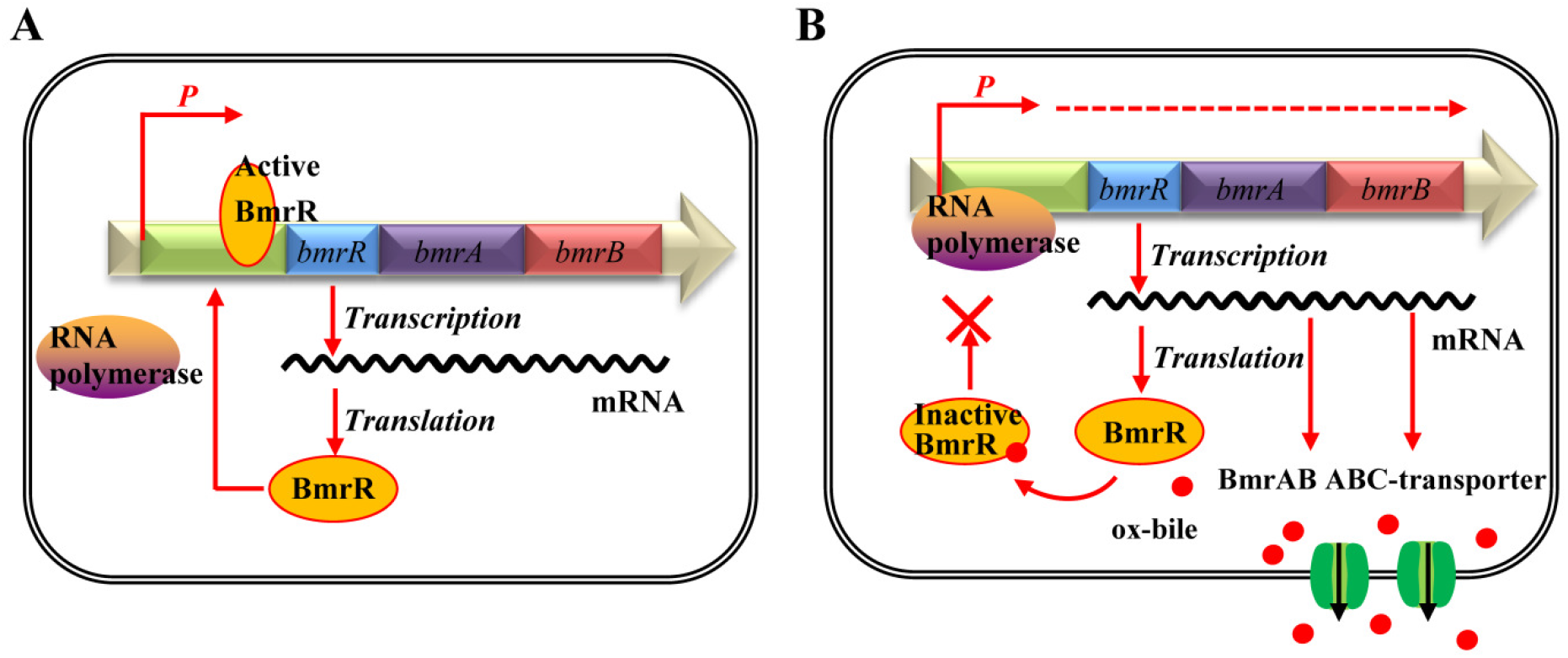
Illustration of the BmrR regulation mechanism. (A)Under the normal growth condition, the active form of BmrR binds to the bmr box and represses transcription of BmrRAB operon. (B) In presence of ox-bile, the DNA binding activity of BmrR is disrupted by ox-bile. This modification will result in the transcription of BmrAB ABC-transporters to pump out ox-bile.

## Material and methods

### Bacterial strains and growth conditions

The bacterial strains and plasmids used in this study are listed in Table S1. *B. longum* BBMN68 was grown in de Man-Rogosa-Sharpe (MRS) broth supplemented with 0.05% (vol/vol) L-cysteine (MRSc) at 37°C anaerobically (5% CO_2_, 5% H_2_ and 90% N_2_). *Lactococcus lactis* NZ9000 was routinely grown at 30°C in M17 medium (Oxoid, Unipath, Basingstoke, UK) containing 0.5% wt/vol glucose (GM17). When necessary, medium was supplemented with 10 μg · ml^−1^ chloramphenicol for *L. lactis.*

### Construction of the recombinant strain *L. lactis* BmrR

Standard PCR was carried out using Q5™ High-Fidelity DNA polymerase following the manufacturer’s instructions (NEB, Beijing, China). The *bmrR* was amplified from genomic DNA of *B. longum* BBMN68 using the primer pair: *bmrR-F* and *bmrR-R* (Table S2). The PCR product digested by *NcoI* and *XbaI* was inserted into the corresponding sites of pNZ8148. Subsequently, the ligation mixture was transformed into *L. lactis* NZ9000 according to previously described procedures (39). The recombinant plasmid pNZBmrR was verified by DNA sequencing and further analyzed with the DNAMAN software package (Lynnon Biosoftware, Vaudreuil, Quebec, Canada). The strain harboring pNZBmrR was designated *L. lactis* BmrR. Meanwhile, a control strain *(L. lactis* NZCK) was constructed by introducing the empty vector pNZ8148 into *L. lactis* NZ9000. Sodium dodecyl sulphate-polyacrylamide gel electrophoresis (SDS-PAGE) analysis was used to investigate the expression of *bmrR* in *L. lactis.*

### Bile stress survival experiment

Overnight cultures of recombinant strains were inoculated into 10 ml of fresh GM17 supplemented with 10 μg · ml^−1^ chloramphenicol (1% inoculums). When cell density reached an OD_600_ of 0.3, nisin was added (final concentration 10 ng · μl^−1^) and further incubated for 2 h at 30°C. Aliquots of 1 mL of culture were collected and suspended in 1ml fresh GM17 medium containing 0.10% wt/vol ox-bile (Sigma, St. Louis, MO, USA). After incubation at 30°C for 1 h, the number of colony-forming units per milliliter (CFU · ml^−1^) was determined by plating 10-fold serial dilutions on GM17 plates containing 10 μg · μl^−1^ chloramphenicol and incubating at 30°C for 16 h. Survival rate were calculated by dividing the number of colony-forming units (CFU) per ml after ox-bile incubation by the value obtained immediately after resuspension. All results were obtained by at least three independent experiments with each performed in triplicate.

### Purification of recombinant BmrR and electrophoretic mobility shift assay (EMSA)

The gene *bmrR* was amplified by PCR using the primer pair *bmrRHis*-F and *bmrRHis*-R listed in Table S2, which introduced a six histidine tag at the C-terminal end of this protein, immediately prior to the stop codon to simplify protein purification by affinity chromatography using a nickel column. The PCR product digested by *NcoI/XbaI* was ligated with pNZ8148 at the corresponding restriction sites, resulting in recombinant plasmid pBmrRHis. This plasmid was then introduced into *L. lactis* NZ9000, and the transformant harboring the correct construct was designated *L. lactis* BmrRHis. The protein BmrR with a C-terminal His tag (designated BmrRHis) was purified with Ni Sepharose 6 Fast Flow media (GE Healthcare, Uppsala, Sweden) according to the manufacturer’s recommendations. Subsequently, purified BmrRHis was concentrated by ultrafiltration (Millipore, 10 kDa cut off; Bedford, MA, USA) and centrifugation at 13,000 × *g* for 30 min at 4°C. The purified BmrRHis was analyzed by SDS-PAGE and protein concentration was estimated using NanoDrop 2000 UV-Vis Spectrophotometer (Thermo Scientific, Wilmington, DE, USA). Purified protein was used immediately or stored at -80°C for subsequent experiments.

EMSA was performed using the LightShift® Chemiluminescent EMSA Kit (Thermo Scientific, Rockford, IL, USA). To obtain biotin 3’ end-labelled probes, two complementary oligonucleotides listed in Table S2 were synthesized and annealed at 95°C for 5 min, with the temperature decreasing by 1°C per minute and thereafter until holding 4°C. EMSA were performed according to the manufacturer’s instructions, and the binding reactions (20 μl) contained 1×binding buffer, 50 ng · μl’^1^ Poly dI-dC, 2.5% (vol/vol) glycerol, 0.05% (vol/vol) NP-40, 20 fmol labeled probe, 5 mM MgCl_2_, and 0.01 ng · μl^−1^ BmrRHis, for 20 min at room temperature. To determine if the inverted repeat (IR) structure of predicted binding site was essential, conserved binding site IR1 (<u>ATTGTTG</u>) was changed to <u>GCCACGA</u>, and IR2 (<u>CAACAAT</u>) was changed to <u>GCCACGA</u>, as shown in Figure 4B. In addition, different concentrations of BmrRHis (0.01 μg · μl^−1^, 0.02μg · μl^−1^ and 0.03μg · μl^−1^) and ox-bile (0.05%, 0.10% and 0.15% wt/vol) were applied to determine the dose effects on the binding activity of BmrR. The subsequent steps were carried out following the manufacturer’s instructions. Chemiluminescent signals of biotinylated probes were captured using a CCD camera imaging system (UVP, Upland, CA, USA).

### Validation of *bmrR* operon by Reverse-transcription PCR

*B. longum* BBMN68 cells were immediately harvested at an OD_600_ of 0.6 by centrifugation at 6000 × *g* for 10 min. Total RNA was isolated using Trizol Reagent (Invitrogen, Carlsbad, CA, USA) according to the manufacturer’s instructions, and digested with RNase-free DNase I (Tiangen, Beijing, China). RNA concentrations were quantified using the NanoDrop 2000 (Thermo Scientific). RNA quality was assessed with the 2100 Bioanalyzer (Agilent Technologies, Amstelveen, Netherlands). Subsequently, reverse transcription was carried out with PrimeScript II 1st strand cDNA synthesis kit (Takara, Beijing, China), with 1 μg of total RNA as the template. Specific primers listed in Table S2 were designed using PRIMER V5 software (PREMIER Biosoft International, Palo Alto, CA). Standard PCR was carried out using Q5™ High-Fidelity DNA polymerase (NEB) with the cDNA as template, RNA as negative control and the genomic DNA of BBMN68 as positive control.

### Functional identification of target gene *bmrA* and *bmrB* by Heterologous Expression

The *bmrA* gene and *bmrB* gene encoding half ABC-transporter were amplified from *B. longum* BBMN68 using primers *bmrA-F/bmrA-R* and bmrB-F/bmrB-R, respectively (Table S2). Then the *bmrA* and *bmrB* gene was co-amplified using *bmrA-F* and *bmrB-R.* All PCR amplicons digested by XbaI and HindIII were inserted into pNZ8147. The ligation mixture was transformed into *L. lactis* NZ9000, resulting in recombinant strain *L. lactis* BmrA, *L. lactis* BmrB and *L. lactis* BmrAB, respectively. Meanwhile, the strain *L. lactis* NZ9000 with the empty vector pNZ8147 was used as control. The recombinant strains were grown in GM17 medium supplemented with 0. 05%, 0.10% or 0.15% wt/vol ox-bile to determine the survival rate. All results were obtained by at least three independent experiments with each performed in triplicate.

### Statistical analysis

Data were analyzed using GraphPad Prism 6 software for Windows (GraphPad Software, Inc., La Jolla, CA, USA). When two groups were compared, an unpaired student t test with Welch’s correction to calculate P values. When three groups or more were compared, one-way ANOVA was used followed by an appropriate post-hoc test.

## Authors and Contributors

Q.X., H.A., Z.Z. and Y.H. designed research; Q.X. and H.A. performed research; Y.Y., J.Y., and G.W. contributed new reagents or analytic tools; Q.X., Z.Z. and Y.H. analyzed the data and wrote the paper. All authors read and approved the final manuscript.

## Acknowledgements

This work was supported by the National Natural Sciences Foundation of China (contract No.21676294) and the International Postdoctoral Exchange Fellowship Program [No. 20150027]. We thank Professor Willem M. de Vos (Wageningen University) for the gift of *L. lactis* NZ9000 and plasmid pNZ8148. We also thank Dr. Kuanqing Liu (UT Southwestern Medical Center) for helpful comments.

## References

1. Harmsen HJ1, Wildeboer-Veloo AC, Raangs GC, Wagendorp AA, Klijn N, Bindels JG, Welling GW. 2000. Analysis of intestinal flora development in breast-fed and formula-fed infants by using molecular identification and detection methods. J Pediatr Gastroenterol Nutr 30: 61–67.

2. Hofmann AF, and Eckmann L. 2006. How bile acids confer gut mucosal protection against bacteria. Proc Natl Acad Sci U S A 103: 4333–4334.

3. Tannock GW. 1999. A fresh look at the intestinal microflora. In: Tannock GW, editor. Probiotics. a critical review. Wymondham, (UK): Horizon Scientific Press.

4. Noriega L, Gueimonde M, Sánchez B, Margolles A, de los Reyes-Gavilán CG. 2004. Effect of the adaptation to high bile salts concentrations on glycosidic activity, survival at low pH and cross-resistance to bile salts in *Bifidobacterium*. Int J Food Microbiol 94: 79–86. doi: 10.1016/j.ijfoodmicro.2004.01.003

5. Sánchez B, Champomier-Vergès MC, Stuer-Lauridsen B, Ruas-Madiedo P, Anglade P, Baraige F, de los Reyes-Gavilán CG, Johansen E, Zagorec M, Margolles A. 2007. Adaptation and response of *Bifidobacterium animalis* subsp. *lactis* to bile: a proteomic and physiological approach. Appl Environ Microbiol 73: 6757–6767. doi: 10.1128/aem.00637-07.

6. Bernstein C, Bernstein H, Payne CM, Beard SE, Schneider J. 1999. Bile salt activation of stress response promoters in *Escherichia coli*. Curr Microbiol 39: 68–72.

7. Grill JP, Perrin S, and Schneider F. 2000. Bile salt toxicity to some bifidobacteria strains: role of conjugated bile salt hydrolase and pH. Can J Microbiol 46: 878–884.

8. Price CE, Reid SJ, Driessen AJ, Abratt VR. 2006. The *Bifidobacterium longum* NCIMB 702259T ctr gene codes for a novel cholate transporter. Appl Environ Microbiol 72: 923–926.

9. Gueimonde M, Garrigues C, van Sinderen D, de los Reyes-Gavilan CG, Margolles A. 2009. Bile-inducible efflux transporter from *Bifidobacterium longum* NCC2705, conferring bile resistance. Appl Environ Microbiol 75: 3153–3160.

10. Margolles A, Florez AB, Moreno JA, van Sinderen D, de los Reyes-Gavilan CG. 2006. Two membrane proteins from *Bifidobacterium breve* UCC2003 constitute an ABC-type multidrug transporter. Microbiology 152: 3497–3505. doi: 10.1099/mic.0.29097-0

11. Ruiz L, Sánchez B, Ruas-Madiedo P, de los Reyes-Gavilán CG, Margolles A. 2007. Cell envelope changes in *Bifidobacterium animalis* ssp. *lactis* as a response to bile. FEMS Microbiol Lett 274: 316–322.

12. Ruiz L, Coute Y, Sanchez B, de los Reyes-Gavilan CG, Sanchez JC, Margolles A. 2009. The cell-envelope proteome of *Bifidobacterium longum* in an *in vitro* bile environment. Microbiology 155: 957–967. doi: 10.1099/mic.0.024273-0

13. Liu Y, An H, Zhang J, Zhou H, Ren F, Hao Y. 2014. Functional role of *tlyC1* encoding a hemolysin-like protein from *Bifidobacterium longum* BBMN68 in bile tolerance. FEMS Microbiol Lett 360: 167–173. doi: 10.1111/1574-6968.12601.

14. An H, Douillard FP, Wang G, Zhai Z, Yang J, Song S, Cui J, Ren F, Luo Y, Zhang B, Hao Y. (2014). Integrated transcriptmic and proteomic analysis of the bile stress response in a centenarian-originated probiotic *Bifidobacterium longum* BBMN68. Mol Cell Proteomics 13: 2558–2572. doi: 10.1074/mcp.M114.039156

15. Yang H, Liu A, Zhang M, Ibrahim SA, Pang Z, Leng X, Ren F. 2009. Oral administration of live *Bifidobacterium* substrains isolated from centenarians enhances intestinal function in mice. Curr Microbiol 59: 439–445. doi: 10.1007/s00284-009-9457-0

16. Yang J, Zhang H, Jiang L, Guo H, Luo, X, Ren F. 2015. *Bifidobacterium longum* BBMN68-specific modulated dendritic cells alleviate allergic responses to bovine beta-lactoglobulin in mice. J Appl Environ Microbiol 119: 1127–1137. doi: 10.1111/jam.12923

17. Wilkinson SP, Grove A. 2006. Ligand-responsive transcriptional regulation by members of the MarR family of winged helix proteins. Curr Issues Mol Biol 8: 51–62.

18. Wagner A, Segler L, Kleinsteuber S, Sawers G, Smidt H, Lechner U. 2013. Regulation of reductive dehalogenase gene transcription in *Dehalococcoides mccartyi*. Philos Trans R Soc Lond B Biol Sci 368, 20120317. doi: 10.1098/rstb.2012.0317

19. Martin RG, Rosner JL. 1995. Binding of purified multiple antibiotic-resistance repressor protein (MarR) to mar operator sequences. Proc Natl Acad Sci U S A 92: 5456–5460.

20. Lomovskaya O, Lewis K, Matin A. 1995. EmrR is a negative regulator of the *Escherichia coli* multidrug resistance pump EmrAB. J Bacteriol 177: 2328–2334.

21. Schielke S, Huebner C, Spatz C, Nägele V, Ackermann N, Frosch M, Kurzai O Schubert-Unkmeir A. 2009. Expression of the meningococcal adhesin NadA iscontrolled by a transcriptional regulator of the MarR family. Mol Microbiol 72: 1054–1067.

22. Perera IC, Grove A. 2010. Molecular mechanisms of ligand-mediated attenuation of DNA binding by MarR family transcriptional regulators. J Mol Cell Biol 2: 243–254.

23. Alekshun MN, Levy SB, Mealy TR, Seaton BA, Head JF. 2001. The crystal structure of MarR, a regulator of multiple antibiotic resistance, at 2.3 A resolution. Nat Struct Biol 8: 710–714. doi: 10.1038/90429

24. Sulavik MC, Gambino LF, Miller PF. 1995. The MarR repressor of the multiple antibiotic resistance (mar) operon in *Escherichia coli*: prototypic member of a family of bacterial regulatory proteins involved in sensing phenolic compounds. Mol Med 1: 436–446.

25. Alekshun MN, Kim YS, Levy SB. 2000. Mutational analysis of MarR, the negative regulator of *marRAB* expression in *Escherichia coli*, suggests thepresence of two regions required for DNA binding. Mol Microbiol 35: 1394–1404.

26. Hong M, Fuangthong M, Helmann JD, Brennan RG. 2005. Structure of an *OhrR-ohrA* operator complex reveals the DNA binding mechanism of the MarR family. Mol Cell 20: 131–141. doi: 10.1016/j.molcel.2005.09.013

27. Reese MG. 2001. Application of a time-delay neural network to promoter annotation in the *Drosophila melanogaster* genome. Comput Chem 26: 51–56.

28. Solovyev V, Salamov A. 2011. Automatic annotation of microbial genomes and metagenomic sequences. Metagenomics and its applications in agriculture, biomedicine and environmental studies, 61–78.

29. Deochand DK, Grove A. 2017. MarR family transcription factors: dynamic variations on a common scaffold. Crit Rev Biochem Mol Biol 52: 595–613. doi: 10.1080/10409238.2017.1344612.

30. Grkovic S, Brown MH, Skurray RA. 2002. Regulation of bacterial drug export systems. Microbiol Mol Biol Rev 66: 671–701.

31. Prouty AM, Brodsky IE, Falkow S, Gunn, JS. 2004. Bile-salt-mediated induction of antimicrobial and bile resistance in *Salmonella typhimurium*. Microbiology 150: 775–783.

32. Zaidi AH, Bakkes PJ, Lubelski J, Agustiandari H, Kuipers OP, Driessen AJ. 2008. The ABC-type multidrug resistance transporter LmrCD is responsible for an extrusion-based mechanism of bile acid resistance in *lactococcus lactis*. J Bacteriol 190: 7357–66. doi: 10.1128/JB.00485-08.

33. Michaux C, Martini C, Hanin A, Auffray Y, Hartke A, Giard JC. 2011. SlyA regulator is involved in bile salts stress response of *Enterococcus faecalis*. FEMS Microbiol Lett 324: 142–6. doi: 10.1111/j.1574-6968.2011.02390.x.

34. Schneider E, Hunke S. 1998. ATP-binding-cassette (ABC) transport systems: functional and structural aspects of the ATP-hydrolyzing subunits/domains. FEMS Microbiol Rev 22: 1–20.

35. Walker JE, Saraste M, Runswick MJ, Gay NJ. 1982. Distantly related sequences in the alpha-and beta-subunits of ATP synthase, myosin, kinases and other ATP-requiring enzymes and a common nucleotide binding fold. EMBO J 1: 945–951.

36. van Veen HW, Margolles A, Muller M, Higgins CF, Konings WN. 2000. The homodimeric ATP-binding cassette transporter LmrA mediates multidrug transport by an alternating two-site (twocylinder engine) mechanism. EMBO J 19, 2503–2514.

37. Chang G, Roth CB. 2001. Structure of MsbA from *E. coli*: a homolog of the multidrug resistance ATP binding cassette (ABC) transporters. Science 293: 1793–1800.

38. Lubelski J, Mazurkiewicz P, van Merkerk R, Konings WN, Driessen AJ. 2004. *ydaG* and *ydbA* of *Lactococcus lactis* encode a heterodimeric ATP-binding cassette-type multidrug transporter. J Biol Chem 279: 34449–34455.

39. de Ruyter PG, Kuipers OP, de Vos WM. 1996. Controlled gene expression systems for *Lactococcus lactis* with the food-grade inducer nisin. Appl Environ Microbiol 62: 3662–3667.

